# PlasmidTron: assembling the cause of phenotypes from NGS data

**DOI:** 10.1101/188920

**Authors:** Andrew J. Page, Alexander Wailan, Yan Shao, Kim Judge, Gordon Dougan, Elizabeth J. Klemm, Nicholas R. Thomson, Jacqueline A. Keane

## Abstract

When defining bacterial populations through whole genome sequencing (WGS) the samples often have detailed associated metadata that relate to disease severity, antimicrobial resistance, or even rare biochemical traits. When comparing these bacterial populations, it is apparent that some of these phenotypes do not follow the phylogeny of the host i.e. they are genetically unlinked to the evolutionary history of the host bacterium. One possible explanation for this phenomenon is that the genes are moving independently between hosts and are likely associated with mobile genetic elements (MGE). However, identifying the element that is associated with these traits can be complex if the starting point is short read WGS data. With the increased use of next generation WGS in routine diagnostics, surveillance and epidemiology a vast amount of short read data is available and these types of associations are relatively unexplored. One way to address this would be to perform assembly *de novo* of the whole genome read data, including its MGEs. However, MGEs are often full of repeats and can lead to fragmented consensus sequences. Deciding which sequence is part of the chromosome, and which is part of a MGE can be ambiguous. We present *PlasmidTron*, which utilises the phenotypic data normally available in bacterial population studies, such as antibiograms, virulence factors, or geographic information, to identify sequences that are likely to represent MGEs linked to the phenotype. Given a set of reads, categorised into cases (showing the phenotype) and controls (phylogenetically related but phenotypically negative), *PlasmidTron* can be used to assemble *de novo* reads from each sample linked by a phenotype. A *k*-mer based analysis is performed to identify reads associated with a phylogenetically unlinked phenotype. These reads are then assembled *de novo* to produce contigs. By utilising *k*-mers and only assembling a fraction of the raw reads, the method is fast and scalable to large datasets. This approach has been tested on plasmids, because of their contribution to important pathogen associated traits, such as AMR, hence the name, but there is no reason why this approach cannot be utilized for any MGE that can move independently through a bacterial population. *PlasmidTron* is written in Python 3 and available under the open source licence GNU GPL3 from https://github.com/sanger-pathogens/plasmidtron.

**DATA SUMMARY:**

1. Source code for *PlasmidTron* is available from Github under the open source licence GNU GPL 3; (url - https://goo.gl/ot6rT5)
2. Simulated raw reads files have been deposited in Figshare; (url - https://doi.org/10.6084/m9.figshare.5406355.vl)
3. *Salmonella enterica* serovar Weltevreden strain VNS10259 is available from GenBank; accession number GCA_001409135.
4. *Salmonella enterica* serovar Typhi strain BL60006 is available from GenBank; accession number GCA_900185485.
5. Accession numbers for all of the Illumina datasets used in this paper are listed in the supplementary tables.

**I/We confirm all supporting data, code and protocols have been provided within the article or through supplementary data files.** ⊠

**IMPACT STATEMENT:** PlasmidTron utilises the phenotypic data normally available in bacterial population studies, such as antibiograms, virulence factors, or geographic information, to identify sequences that are likely to represent MGEs linked to the phenotype.

## INTRODUCTION

When defining bacterial populations through whole genome sequencing (WGS) the samples often have detailed associated metadata that relate to disease severity, antimicrobial resistance, or even rare biochemical traits. When comparing these bacterial populations, it is apparent that some of these phenotypes do not follow the phylogeny of the host i.e. they are genetically unlinked to the evolutionary history of the host bacterium. One possible explanation for this phenomenon is that the genes are moving independently between hosts and are likely associated with mobile genetic elements (MGE). However, identifying the element that is associated with these traits can be complex if the starting point is short read WGS data. With the increased use of next generation WGS in routine diagnostics, surveillance and epidemiology a vast amount of short read data is available and these types of associations are relatively unexplored. One way to address this would be to perform assembly *de novo* of the whole genome read data, including its MGEs. However, MGEs are often full of repeats and can lead to fragmented consensus sequences. Deciding which sequence is part of the chromosome, and which is part of a MGE can be ambiguous (1).

A number of recent methods have been developed to address the problem of assembling some of these MGEs, from NGS data (1). *plasmidSPAdes* (2) detects plasmids by analysing the coverage of assembled contigs to separate out chromosomes from plasmid like sequences. By filtering the dataset, a higher quality assembly is possible. However, if the copy number of the plasmids are similar to the chromosome, it is not possible to separate out plasmids. *Unicycler* (3) is a hybrid assembler which can combine short and long read data to produce fully circularised chromosomes and plasmids. It essentially fixes many of the deficiencies of *SPAdes* (4) and fine tunes it for assembling bacteria. *Recycler* (5) takes an assembly graph and aligned reads to search for cycles in the graph which may correspond to plasmids. The method is only partially implemented with substantial work required on the researcher’s part to generate input files in the correct formats. It is shown to work well on small simple plasmids, however it does not scale to larger more complex plasmids. All of these software applications utilise *SPAdes* within their methods, work on a single sample at a time, and require no *a priori* knowledge about the samples themselves.

We present *PlasmidTron,* which utilises the phenotypic data normally available in bacterial population studies, such as antibiograms, virulence factors, or geographic information, to identify sequences that are likely to represent MGEs linked to the phenotype. Given a set of reads, categorised into cases (showing the phenotype) and controls (phylogenetically related but phenotypically negative), *PlasmidTron* can be used to assemble *de novo* reads from each sample linked by a phenotype. A *k-mer* based analysis is performed to identify reads associated with a phylogenetically unlinked phenotype. These reads are then assembled *de novo* to produce contigs. By utilising *k*-mers and only assembling a fraction of the raw reads, the method is fast and scalable to large datasets. This approach has been tested on plasmids, because of their contribution to important pathogen associated traits, such as AMR, hence the name, but there is no reason why this approach cannot be utilized for any MGE that can move independently through a bacterial population. The method is tested on simulated and real datasets, compared to other methods, and the results are validated with long read sequencing. *PlasmidTron* is a command-line tool, is written in Python 3 and is available under the open source licence GNU GPL3 from https://goo.gl/ot6rT5.

## METHOD

*PlasmidTron* takes two spreadsheets as input, one containing paired ended reads in FASTQ format for samples displaying the phenotype (cases), the other containing FASTA or FASTQ files for samples not displaying the phenotype (controls). The full method is shown in Figure 1. A *k*-mer analysis of each of the samples is performed using KMC (syntax versions v2.3.0 or v3.0.0) (6,7) to produce databases of *k*-mer counts, *k*-mers occurring less than 5 times are excluded by default since assembly is more error prone below this level of coverage. A union is taken of the cases *k-mer* databases to produce a new database of all *k*-mers ever seen in any of the trait samples, and similarly for the controls. The two sets are then subtracted from each other, leaving only *k*-mers uniquely present in the cases dataset. The raw reads, plus their mates, which match these unique *k-*mers are extracted from each sample where each read must be covered by a defined percentage of *k*-mers. Each set of reads is assembled *de novo* with SPAdes. The assembly contigs are filtered to remove small contigs (default 300 bases), and low coverage contigs (below 10X). This is because a single erroneous *k-mer* can draw in reads on either side equating to approximately the fragment size of the library. The resulting sequences can be fragmented so a second scaffolding step is undertaken. A k-mer database is generated for each assembly and the raw reads, plus their mates, are extracted for a second assembly with *SPAdes.* This allows for gaps of up to twice the fragment size to be closed. A final filtering step of the assembled sequences is performed, as previously described. An assembly in FASTA format is created for each of the trait samples, along with a plot of the shared *k*-mers in each sample, indicating the level of identity between samples. Parallelisation support is provided by GNU parallel (8).

**Figure 1:**
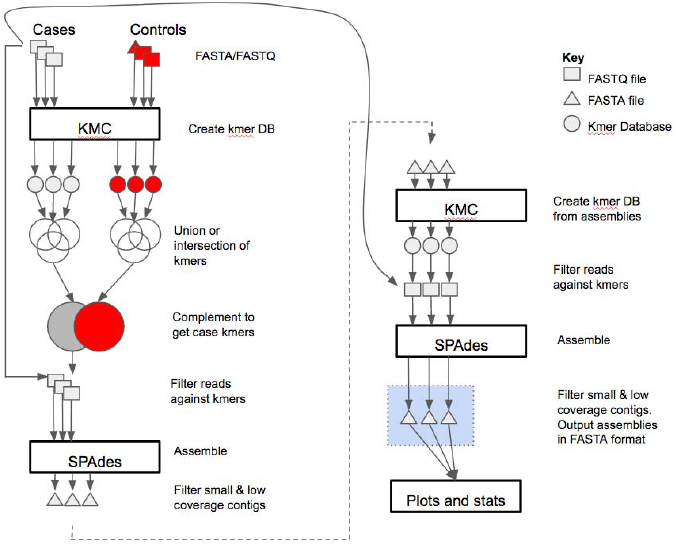
The PlasmidTron algorithm. FASTQfiles are denoted as squares, FASTA files as triangles and *k*-mer databases as circles.

## RESULTS

To evaluate the effectiveness of *PlasmidTron* three datasets were used including: 1) simulated reads to show the impact of copy number variation in identifying plasmids, 2) the effectiveness of different methods in recalling plasmid type sequences on real world data, and 3) identification of a novel AMR plasmid with subsequent validation using long read sequencing. All experiments were performed using the Wellcome Trust Sanger Institute compute infrastructure, running Ubuntu 12.04.

### IMPACT OF COPY NUMBER VARIATION

Simulated reads were generated to show the impact of copy number variation compared to other methods. A trivial set of simulated perfect reads was generated. A reference genome, which was sequenced using the PacBio RSII for *Salmonella enterica* serovar Weltevreden *(S.* Weltevreden) (accession number GCA_001409135), was shredded using FASTAQ (v3.15.0) (https://github.com/sanger-pathogens/fastaq) to generate perfect paired-ended reads with a read length of 125 bases and a mean fragment size of 400 bases. The reference contains a single chromosome (5,062,936 bases) and single plasmid (98,756 bases), where the chromosome depth of coverage was fixed at 30X, and the plasmid depth of coverage was varied from 1 to 60X in steps of 2. The break point for the plasmid was varied, in steps of 500 bases, to simulate a circular genome.

The results of *PlasmidTron* (v0.3.5) were compared to 4 other methods, *recycler* (v0.6), *Unicycler* (vO.4.0), *SPAdes* (v3.10.0), and *plasmidSPAdes* (v3.10.0). *SPAdes* (v3.10.0) was used as the assembler for each of these methods, *recycler* required pre-processing steps using *bwa* (v0.7.12) (9) and *samtools* (vO.1.19) (10). *SPAdes* and *Unicycler* are not dedicated plasmid assemblies and are agnostic to the underlying structures being sequenced, however they provide a good baseline for what is possible, though the final plasmid sequences are contained in a large collection of chromosome sequences. *plasmidSPAdes* and *PlasmidTron* are dedicated plasmid assemblers, and *recycler* is post assembly plasmid analysis tool, with each employing a fundamentally different analysis strategy.

Each resulting assembly was measured based on the percentage of plasmid assembled, how fragmented the plasmid was, and the proportion of non-plasmid bases to plasmid bases (signal to noise ratio). The assemblies are blasted (v.2.6.0) (11) against the expected plasmid sequence, with an e-value of 0.0001. Blast hits of less than 200 bases long or less than 90% identity were excluded. *Recycler* identified no plasmids on the real or simulated data, which appears to be due to the large complex size of the plasmid.

Figure 2 shows that as the copy number of the plasmid in the input reads changes, the percentage of the plasmid recovered changes. *plasmidSPAdes* only begins to start identifying plasmid sequences at 40X, recovering the full plasmid sequence. Below this level the plasmid copy number is too similar to the chromosome coverage so the algorithm filters it out. The *SPAdes* and *Unicycler* assemblers identify all of the plasmid sequence with less than 10X coverage, however the plasmid sequences are fragmented and makes up only ∼1.9% of the final assembly as show in Figure 3. *PlasmidTron* requires slightly more coverage (16X) to generate an assembly which covers the full plasmid sequence. At 16X more than 90% of the resulting assembly contains plasmid sequences, increasing to 100% at 40X.

**Figure 2:**
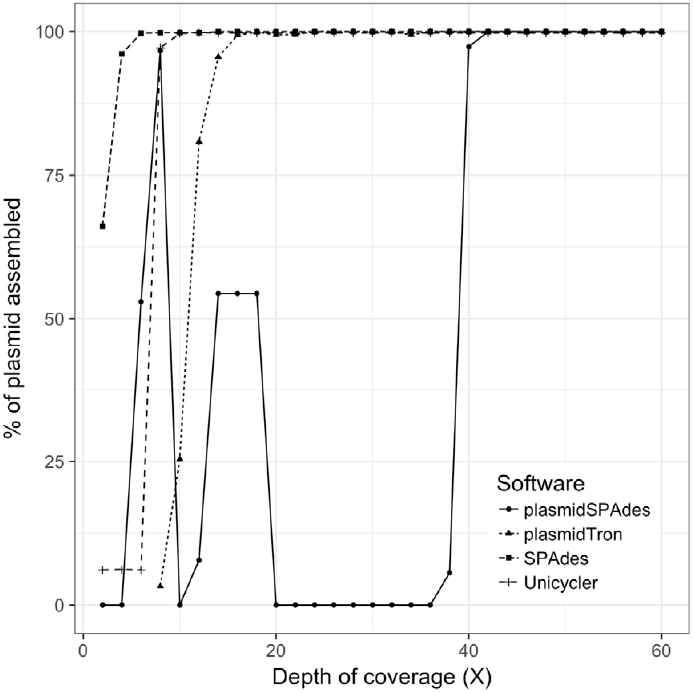
The percentage of the plasmid sequence which was assembled with different software applications as the depth of coverage of a plasmid increases in the raw data.

**Figure 3:**
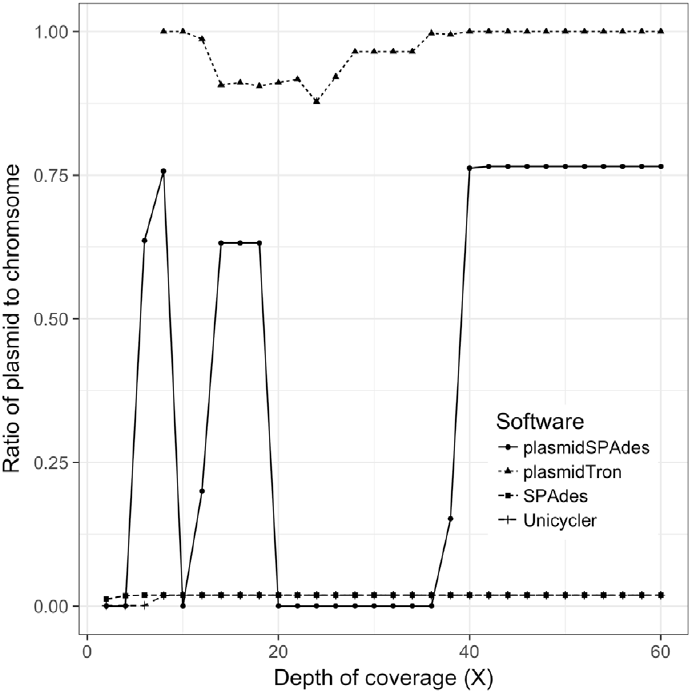
The ratio of the plasmid sequence to the chromosome sequence in the final assembly produced by each software application as the depth of coverage of the plasmid increases in the raw reads. This is akin to the signal to noise ratio.

### RECOVERY OF TYPING SEQUENCE

A real dataset of 114 isolates of S. Weltevreden, was sequencing using Illumina as described in (12). The samples are clonal, with most sharing a similar plasmid, although the payload of the plasmid itself varies greatly. To get a baseline for what plasmids are present in the input dataset, all of the samples were compared to the PlasmidFinder (13) database (retrieved 2017-07-25) using Ariba (v2.10.0) (14), providing the Incompatibility group. PlasmidFinder identifed one plasmid group, IncFlls, as present in 89.5% of samples. *plasmidSPAdes, Unicycler, SPAdes, Recycler* and *PlasmidTron* were provided with the dataset and the results were searched for the IncFIl_s_ sequence using blastn, with full details listed in Supplementary Table 1. In 4 cases PlasmidFinder failed to identify the sequence, when it was found by 2 or more other applications. *SPAdes* and *Unicycler* identify the sequence in 88.6% and 87.7% of samples. *plasmidSPAdes* identifies the plasmid sequence in just 8.8% of samples. Recycler failed to identify the plasmid sequence in any sample. *PlasmidTron* identified the plasmid sequence in 87.7% of cases where the chromosome sequence of S. Weltevreden strain VNS10259 was used as the control, giving identical results to *Unicycler.* The benefit though over *Unicycler* is that the majority of the assembled sequences correspond only to the plasmid.

### OUTBREAK AMR

*PlasmidTron* was used to analyse an outbreak of 87 *Salmonella enterica* serovar Typhi (S. Typhi) samples with a resistance profile which had not been previously observed in the haplotype (H58,4.3.1)(15). Further analysis using PlasmidFinder, as previously described, indicated that the antibiotic resistance may reside on an IncY plasmid, a plasmid type which had not been associated before with this haplotype. The chromosomes of 6 complete reference genomes for S. Typhi were used as the controls (accessions GCA_000195995, GCA_000007545, GCA_001157245, GCA_000245535, GCA_001302605, GCA_000385905) for *PlasmidTron,* and 87 Illumina sequenced outbreak samples were used as cases (Supplementary Table 2). For each outbreak sample, *PlasmidTron* identified similar sequences, split over 4-5 contigs. One contig carried the IncY sequence and a second carried AMR genes. Subsequent resequencing of 1 sample (ERS1670682) using long read technology (Oxford Nanopore MinION), revealed that these 4 sequences comprised a single plasmid (accession number GCA_900185485.1), which was identical in all of the outbreak strains. The sequences generated by *PlasmidTron* recovered an average of 96% of the plasmid sequence. The fragmentation (mean 4.6) of the plasmid in the Illumina sequenced samples was due to repeats which could not be resolved with short read sequencing. Overall 65% of the sequences in the resulting assemblies were part of the plasmid sequence, with the remainder resulting from a phage recombination in the main chromosome. This indicates the power of *PlasmidTron* to rapidly, accurately and cost effectively extract sequences of clinical importance from short reads alone.

## CONCLUSION

We can utilise the wealth of phenotypic data usually generated for bacterial population studies, be it routine diagnostics, surveillance or outbreak investigation, to reconstruct plasmids responsible for a particular phenotype. Rather than just identifying that an AMR or virulence gene exists in a sample, *PlasmidTron* can reconstruct all of the sequences of the plasmid it is carried on, providing more insight into the underlying mechanisms. We demonstrated with simulated and real sequences that *PlasmidTron* more accurately reconstructs large plasmids compared to other methods. We present the results of a real outbreak of S. Typhi where *PlasmidTron* was used to identify the plasmid sequence carrying a novel AMR resistance profile, not previously described in S. Typhi H58/4.3.1, and validated the results using long read sequencing. Whilst plasmid assembly remains difficult with short reads, *PlasmidTron* allows for phenotypic data to be utilised to greatly reduce the complexity of the challenge.

## AUTHOR STATEMENTS

This work was supported by the Wellcome Trust (grant WT 098051). Thanks to Martin Hunt Kathryn Holt and Ryan Wick for their helpful discussions and assistance with this work. Thanks also to Daryl Domman for naming the software.

## ABBREVIATIONS

MGE: mobile genetic element

WGS: whole genome sequencing

AMR: anti-microbial resistance

